# InterLig: a fast and accurate software for ligand-based virtual screening

**DOI:** 10.1101/544874

**Authors:** Claudio Mirabello, Björn Wallnery

## Abstract

In the past few years, drug discovery processes have been relying more and more on computational methods to sift out the most promising molecules before time and resources are spent to test them in experimental settings. Whenever the protein target of a given disease is not known, it becomes fundamental to have accurate methods for ligand-based Virtual Screening, which compare known active molecules against vast libraries of candidate compounds. Recently, 3D-based similarity methods have been developed that are capable of scaffold-hopping and to superimpose matching molecules. Here, we present InterLig, a new method for the comparison and superposition of small molecules based on 3D, topologically-independent alignments of atoms. We test InterLig on a standard benchmark and show that it compares favorably to the best currently available 3D methods.

InterLig is open source and is available to everyone at: http://wallnerlab.org/interlig.

## 1. Introduction

Virtual Screening (VS) is a computational technique for the discovery of new, biologically active drug molecules. The idea behind VS is to analyze vast databases of libraries of untested compounds with in-silico methods that should sift out the most promising leads before these are tested in experimental settings. Given the costs associated with laboratory experiments, it is no surprise that huge efforts are being put in developing more accurate methods for VS, so that fewer resources are wasted in pursuing potential dead ends. There are two main approaches to VS, based on structure or ligand. In structure-based VS (SBVS) candidate ligands are docked on the structure of a known receptor [1], while in ligand-based VS (LBVS) the similarity of an active ligand is used to expand the number of potential canditate ligands. The SBVS explicitly model the ligand-receptor interaction enable potentially better results [9]. However, when the structure of the receptor is unknown or not accurate enough LBVS can still be used.

LBVS methods are based on the assumption that structurally similar compounds have a higher chance of binding to the same receptor [2]. The structural similarity can be calculated by comparing 2D fingerprints of compounds or by more accurate 3D structural alignments [11, 5] that also allow scaffold hopping [6, 10] and provide starting points for 3D docking.

Most methods using 3D are shape-based, representing molecules as a mixture of gaussians and structure comparisons as overlaps between them [3, 12, 11], more recently more detailed atom-level comparisons have shown promising results [5].

In this study, we present InterLig, an open-source software for 3D-based LBVS. InterLig uses a simulated annealing-based procedure to map sets of atoms from two molecules in a topologically-independent fashion, which makes it particularly suited for scaffold hopping. The simulated annealing procedure makes InterLig especially fast, making it possible to compare tens of thousand of molecules within minutes (see Table S5 in the supplementary). Moreover, along with the similarity score, a p-value is calculated to assess the statistical significance. InterLig is benchmarked against two state-of-the-art softwares for 3D LBVS and outperforms both according to several standard performance measures.

## 2. Datasets

InterLig is benchmarked against DUD-E (Database of Useful Decoys: Enhanced) [7], a standard benchmark for VS softwares. DUD-E contains 22,886 active ligands against 102 protein targets and 50 times more inactive decoys with similar physico-chemical properties but dissimilar 2D topology.

To account for flexible molecules, multiple conformers of DUD-E ligands and decoys are generated using OMEGA (OpenEye Inc.) [4] with the “strict” flag set to false and minimum RMSD between two conformers set to 2Å. Artpproximately 300K additional ligands and 8M additional decoys are generated this way.

## 3. Results and Discussion

InterLig is based on the InterComp algorithm that we recently developed and successfully applied to the comparison of protein interfaces [8]. It is capable of performing topologically-independent alignments of sets of atoms in a 3D space while taking into account both the relative position and the chemical similarity of the aligned atoms. The core of the adapted algorithm in InterLig is identical to InterComp, the difference is only in two parameters involving a cutoff distance, and the trade-off between coordinate and chemical similarity. In addition, to account for the differences in similarity measure the statistical significance of a hit also had to be recalculated. Because the similarity measure used by InterLig is dependent on the size of the compounds, and smaller ligands have a higher probability to obtaining a high score by chance (Fig. S2), the significance (p-value) of a score is calculated by fitting an extreme value distribution to scores between non-related compounds of different sizes in Fig. S2. For more details on the algorithm and the statistical significance, see Supplementary information.

InterLig is benchmarked using standard performance measures for VS (see Supplementary information) against LS-align [5] and LIGSIFT [11], two state-of-the-art softwares for 3D LBVS. LS-align has, to the best of our knowledge, the best performance on the DUD-E benchmark, while LIGSIFT had the best performance on the older DUD set. To ensure a fair comparison each software was run using the default parameters both the DUD-E benchmark and the multiple conformer set, if more than one similarity measure is reported, the measure that showed the highest performance was used. LS-align has a “flexible” option to generate its own set of conformers, however when benchmarked it showed better performance in “rigid” mode with the multiple conformers generated as above (see Table S4). As in previous studies [13], whenever multiple ligands or decoys have the same ID (including multiple conformers), only the most similar to the seed ligand (highest ranked) is used in the analysis.

In the test on the regular DUD-E dataset, InterLig has signficantly larger (P*<*0.05) AUC and enrichment factors across all top rank percentages compared to both LIGSIFT and LS-align, see Table 1. The performance metrics for LS-align are actually slightly better than those reported in the original publication [5], most likely because how multiple compounds with the same ID are treated. Looking per target, InterLig has a higher AUC compared to LS-align for 60 and LIGSIFT for 76 (out of 102) targets, see Fig. S3(a).

**Tab. 1:** Average Enrichment Factors (EF) for different percentage of top hits and Average Area Under the Curve (AUC) on the DUD-E dataset. The highest values for each column are highlighted in bold. The significance of the improvement between InterLig and the other methods is reported for each measure in brackets (Wilcoxon signed-rank test p-value).

| | $EF^{1\%}$ | $EF^{5\%}$ | $EF^{10\%}$ | AUC |
| --- | --- | --- | --- | --- |
| LIGSIFT | 16.88 ($10^{-8}$) | 6.16 ($10^{-6}$) | 3.95 ($10^{-6}$) | 0.71 ($10^{-7}$) |
| LS-align | 20.70 (0.05) | 7.19 (0.03) | 4.44 (0.01) | 0.75 (0.009) |
| InterLig | **22.28** | **7.63** | **4.77** | **0.76** |

For the multiple conformers set, InterLig is significantly better compared to LIGSIFT and LS-align on all performance metrics, except for EF^1%^ on which it does have a higher EF but not in a significant way, see Table 2. Tables with detailed target-by-target results are available for regular in Table S2 and for multiple conformers in Table S3. The results for multiple conformers is overall slightly better compared single conformers, indicating that it might be worth spending some additional time generating conformers to achieve optimal performance (Fig. S4). However, the difference is not huge and if speed is of essence it is almost as good to not generate the conformers.

**Tab. 2:** Average Enrichment Factors (EF) for different percentage of top hits and Average Area Under the Curve (AUC) on the DUD-E multiple conformer dataset. The highest values for each column are highlighted in bold. The significance of the improvement between InterLig and the other methods is reported for each measure in brackets (Wilcoxon signed-rank test p-value).

| | $EF^{1\%}$ | $EF^{5\%}$ | $EF^{10\%}$ | AUC |
| --- | --- | --- | --- | --- |
| LIGSIFT | 22.18 (0.1) | 7.48 (0.019) | 4.63 (0.03) | 0.75 (0.003) |
| LS-align | 22.77 (0.16) | 7.50 (0.002) | 4.63 (0.005) | 0.75 (0.001) |
| InterLig | **23.13** | **8.07** | **4.91** | **0.78** |
| InterLig + LS-align | **25.12** | **8.72** | **5.27** | **0.79** |

It was further noted that the per target performance for InterLig and LS-align were quite different, see Fig. S3. Thus, there should be potential to combine the two approaches to achieve even higher performance. To test this hypothesis, a combination of InterLig and LS-align was constructed by multiplying the reported p-values and resort the hits. Indeed, the combination InterLig+LS-align is superior to both individual methods, demonstrating that the results from the two methods are complementary, see Table 2.

