## Supplementary information for "InterLig: a fast and accurate software for ligand-based virtual screening"

### Algorithm

InterLig is based on the InterComp algorithm that we have recently developed and successfully applied to the comparison of protein interfaces [1]. We use a modified version of a stochastic method for molecular structure matching [2] to generate an optimal mapping between heavy atoms from two molecules with no topologic constraints. The objective function is calculated by comparing atom-to-atom distance maps of the two molecules $a$ and $b$ with length $L_a = N$ and $L_b = M$ $(N \leq M)$. The rows and columns of the distance maps are permutated (Fig. S1(b)) to maximize a linear combination of two similarity functions.

The first similarity function is based on a structural comparison:

$$strdist(D_a, D_b) = \frac{1}{N^2} \sum_{x=1}^{N} \sum_{y=1}^{N} \frac{1}{1 + (\delta_{xy}/d_0)^2} \quad (1)$$

where $\delta_{xy}$ is the absolute difference of the element $(x, y)$ in $D_a$ and $D_a$ (Fig. S1(a)). The $M - N$ atoms (for $x > N$ and $y > N$) from $b$ are not included in the alignment and are thus excluded from the similarity score. Contrarily to InterComp, here the distance values in the distance maps are squared and the default value for $d_0$ is 0.6

To also consider the chemical compatibility of the molecules, a second similarity function is based on atom similarity:

$$seqdist(S_a, S_b) = \frac{1}{N} \sum_{z=1}^{N} I(s_z^a, s_z^b) \quad (2)$$

In this case, a simple identity matrix $I$ is used that gives a score of 1 if two heavy atoms are of the same type and 0 otherwise. The atom alphabet is made of 23 different symbols and is shown in Table S1. Hydrogens (H) are ignored by default in InterLig.

The two similarity functions are combined following the formula:

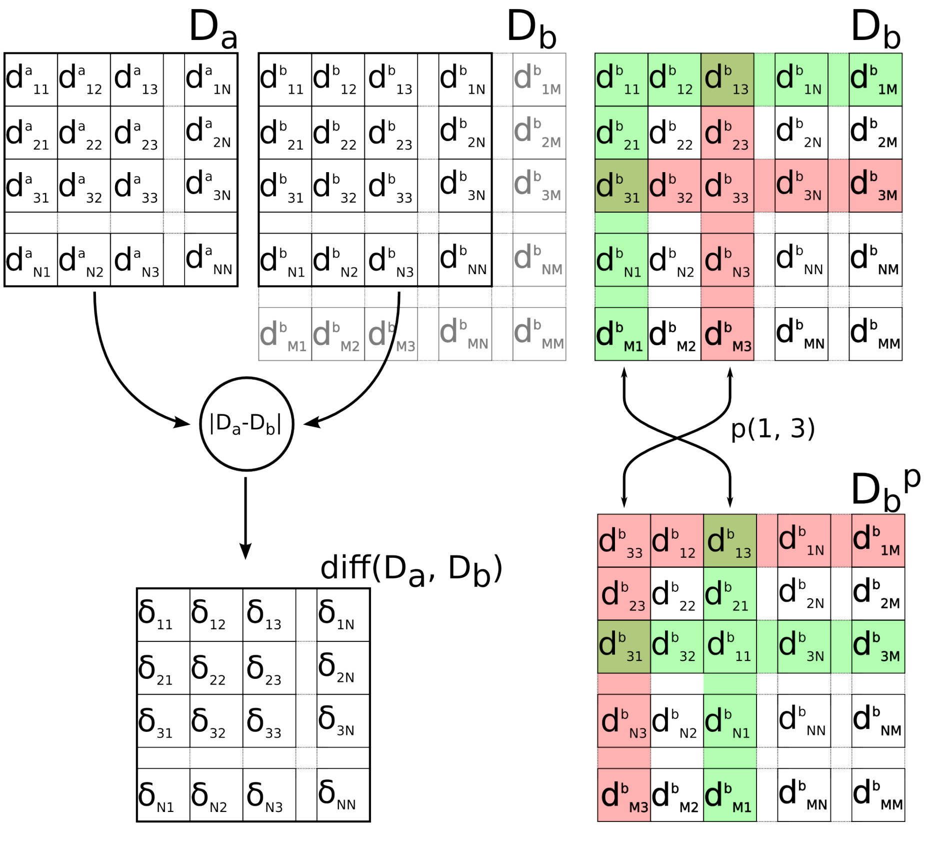


(a) (b)

Fig. S1: (a): The matrix of deltas $\mathrm{diff}(D_a, D_b)$ is the absolute difference, calculated element by element, between the first $N$ rows/columns in the distance matrix $D_a$ and the matrix $D_b$. (b) Permutation of two rows/columns in $D_b$ to form a new candidate mapping $D_b^p$

$$\begin{aligned} opt(p) = \underset{p \in P(M,N)}{\arg\max}\; & W_{str}\, strdist(D_a, D_b^p) + \\ & + (1 - W_{str})\, seqdist(S_a, S_b^p) \end{aligned} \tag{3}$$

where the default value of the weight of the structural component, $W_{str}$, is 0.25.

The default $d_0$ and $W_{str}$ were decided an older release of the dataset used for benchmarking with different ligand and decoys [3].

| | | | | | | |
|---|---|---|---|---|---|---|
| F | F.3 | | | | | |
| Cl | Br | H | | | | |
| S.2 | S.3 | S.02 | | | | |
| O.2 | O.3 | O.co2 | | | | |
| C.1 | C.2 | C.3 | C.ar | | | |
| N.1 | N.2 | N.3 | N.4 | N.am | N.ar | N.pl3 |

Tab. S1: Types of atoms in Virtual Screening of small molecules.

### Assessing statistical significance

In order to assess the statistical significance of potential hits we generated similarity score distributions for alignments of random ligand pairs. The random pairs were obtained by running all-against-all comparison between the seed ligands in the DUD-E set, and pairing up those with lowest similarity scores. The final set of 1.5 million random alignment were then obtained by aligning random molecules from the decoy sets of the previously paired-up seed ligands. The statistical analysis of the distributions of random similarity scores show that they depend on the size of the smaller of the two ligands in the aligment (Fig. S2).

### Performance measures

For each of the 102 targets, we run a LBVS procedure by matching the seed ligand to the ligands and the decoys, and the matched compounds are ranked by their similarity to the seed ligand. In order to assess how well InterLig can sift out biologically active compounds compared to other softwares, we calculate the Enrichment Factor (EF) of $N_{selected}$ molecules at the top $x\%$ of a ranked list of $N_{total}$ compounds:

$$EF^{x\%} = \frac{True\ Positives^{x\%}/N^{x\%}_{selected}}{Total\ Positives/N_{total}} \tag{4}$$

Where $x$ is 1, 5 or 10 (enrichment at top 1%, 5% and 10%). We also calculate the Hit Rate (HR):

$$HR^{x\%} = \frac{EF^{x\%}_{actual}}{EF^{x\%}_{ideal}} \tag{5}$$

Where $EF^{x\%}_{ideal}$ is the best possible Enrichment at the top $x\%$ of the ranking.

We also calculate the Area Under the Curve (AUC) from the Receiving Operating Characteristic (ROC) curve obtained by plotting the True Positive Rate (TPR) against the False Positive Rate (FPR) for each element of the ranked list of molecules.

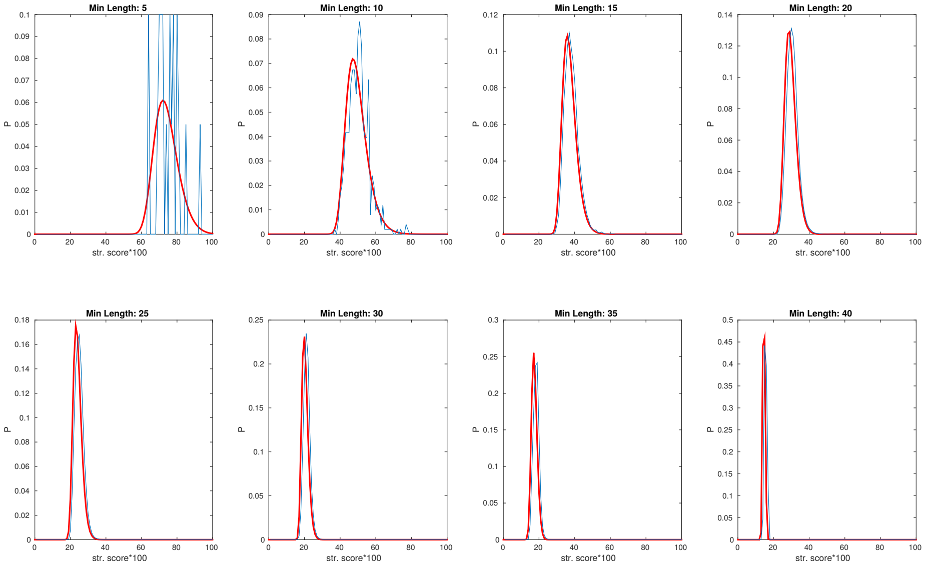


Fig. S2: The similarity score calculated by InterLig is mostly dependent on the size of the smaller of the two molecules being compared. Here we plot the density distributions of InterLig similarity scores for random pairs of non-related compounds depending on the size (number of heavy atoms) of the smaller molecule in the comparison. The empirical probability density is shown in blue, the fitted extreme value function is shown in red.

| Target | $EF^{1\%}$ | $EF^{5\%}$ | $EF^{10\%}$ | $HR^{1\%}$ | $HR^{5\%}$ | $HR^{10\%}$ | AUC |
|---|---|---|---|---|---|---|---|
| aa2ar | 37.53 | 10.04 | 5.95 | 56.56 | 50.20 | 59.50 | 0.782 |
| abl1 | 13.22 | 3.63 | 2.53 | 22.02 | 18.14 | 25.30 | 0.69 |
| ace | 32.70 | 11.14 | 6.99 | 53.79 | 55.70 | 69.90 | 0.909 |
| aces | 15.22 | 7.60 | 4.39 | 25.84 | 38.00 | 43.90 | 0.661 |
| ada | 31.42 | 11.19 | 6.35 | 52.73 | 55.92 | 63.50 | 0.833 |
| ada17 | 27.47 | 9.10 | 5.30 | 40.21 | 45.50 | 53.00 | 0.801 |
| adrb1 | 20.23 | 8.34 | 6.15 | 31.06 | 41.68 | 61.50 | 0.858 |
| adrb2 | 5.20 | 2.51 | 2.12 | 7.89 | 12.54 | 21.20 | 0.665 |
| akt1 | 9.57 | 4.30 | 3.75 | 16.77 | 21.50 | 37.50 | 0.75 |
| akt2 | 16.26 | 4.11 | 3.33 | 27.14 | 20.52 | 33.30 | 0.762 |
| aldr | 31.92 | 9.93 | 5.73 | 55.45 | 49.67 | 57.30 | 0.854 |
| ampc | 28.97 | 7.50 | 4.58 | 48.28 | 37.50 | 45.80 | 0.716 |
| andr | 21.58 | 7.21 | 4.72 | 39.73 | 36.07 | 47.20 | 0.763 |
| aofb | 1.64 | 1.80 | 1.64 | 2.85 | 9.00 | 16.40 | 0.648 |
| bace1 | 8.11 | 2.83 | 2.37 | 12.50 | 14.15 | 23.70 | 0.675 |
| braf | 24.99 | 7.76 | 4.74 | 37.63 | 38.82 | 47.40 | 0.781 |
| cah2 | 0.41 | 2.24 | 6.65 | 0.64 | 11.20 | 66.50 | 0.89 |
| casp3 | 7.53 | 4.12 | 2.91 | 13.76 | 20.62 | 29.10 | 0.618 |
| cdk2 | 5.28 | 2.70 | 2.26 | 8.84 | 13.50 | 22.60 | 0.648 |

| Target | $EF^{1\%}$ | $EF^{5\%}$ | $EF^{10\%}$ | $HR^{1\%}$ | $HR^{5\%}$ | $HR^{10\%}$ | AUC |
|---|---|---|---|---|---|---|---|
| comt | 51.08 | 16.13 | 9.51 | 53.85 | 80.45 | 95.10 | 0.9641 |
| cp2c9 | 0.00 | 0.50 | 1.08 | 0.00 | 2.50 | 10.81 | 0.5467 |
| cp3a4 | 4.11 | 3.41 | 2.82 | 5.84 | 17.04 | 28.20 | 0.673 |
| csf1r | 27.73 | 9.63 | 5.78 | 37.39 | 48.20 | 57.80 | 0.805 |
| cxcr4 | 38.01 | 9.02 | 5.99 | 44.12 | 45.03 | 59.96 | 0.852 |
| def | 25.48 | 10.39 | 6.37 | 44.83 | 51.98 | 63.70 | 0.867 |
| dhi1 | 11.20 | 4.72 | 3.27 | 18.79 | 23.61 | 32.70 | 0.648 |
| dpp4 | 17.47 | 5.67 | 3.62 | 22.46 | 28.35 | 36.20 | 0.745 |
| drd3 | 6.88 | 2.04 | 1.33 | 9.57 | 10.20 | 13.30 | 0.602 |
| dyr | 27.71 | 12.55 | 7.71 | 36.78 | 62.75 | 77.10 | 0.916 |
| egfr | 26.54 | 11.26 | 6.49 | 40.45 | 56.30 | 64.90 | 0.876 |
| esr1 | 45.01 | 12.43 | 7.23 | 81.91 | 62.12 | 72.30 | 0.895 |
| esr2 | 23.22 | 10.20 | 6.16 | 41.47 | 50.97 | 61.60 | 0.862 |
| fa10 | 9.29 | 3.61 | 2.83 | 24.26 | 18.05 | 28.30 | 0.684 |
| fa7 | 8.72 | 9.12 | 6.23 | 15.63 | 45.60 | 62.30 | 0.891 |
| fabp4 | 33.99 | 6.80 | 5.10 | 57.14 | 34.05 | 51.05 | 0.79 |
| fak1 | 33.31 | 17.03 | 9.30 | 61.12 | 84.98 | 93.00 | 0.9635 |
| fgfr1 | 0.68 | 0.85 | 1.01 | 20.00 | 25.00 | 29.71 | 0.477 |
| fkb1a | 12.64 | 4.68 | 3.24 | 23.74 | 23.44 | 32.40 | 0.749 |
| fnta | 20.11 | 7.77 | 5.17 | 22.88 | 38.85 | 51.70 | 0.799 |
| fpps | 90.66 | 19.78 | 9.88 | 90.59 | 98.80 | 98.80 | 0.998 |
| gcr | 26.05 | 7.52 | 4.61 | 44.09 | 37.58 | 46.10 | 0.775 |
| glcm | 3.66 | 2.22 | 2.22 | 5.13 | 11.12 | 22.18 | 0.697 |
| gria2 | 39.84 | 10.62 | 6.01 | 52.50 | 53.15 | 60.10 | 0.805 |
| grik1 | 13.96 | 5.75 | 4.16 | 21.21 | 28.72 | 41.60 | 0.724 |
| hdac2 | 12.41 | 3.57 | 2.49 | 21.90 | 17.84 | 24.90 | 0.631 |
| hdac8 | 5.89 | 4.23 | 3.35 | 9.43 | 21.15 | 33.50 | 0.7208 |
| hivint | 4.03 | 3.20 | 2.70 | 5.98 | 15.99 | 26.97 | 0.753 |
| hivpr | 28.38 | 9.07 | 5.65 | 41.99 | 45.35 | 56.50 | 0.820 |
| hivrt | 8.00 | 3.85 | 2.93 | 14.07 | 19.25 | 29.30 | 0.661 |
| hmdh | 47.13 | 12.46 | 7.06 | 89.89 | 62.36 | 70.60 | 0.899 |
| hs90a | 11.45 | 5.45 | 4.09 | 20.41 | 27.28 | 40.94 | 0.795 |
| hxk4 | 29.27 | 8.27 | 5.65 | 56.25 | 41.29 | 56.50 | 0.845 |
| igf1r | 16.28 | 4.86 | 3.18 | 25.53 | 24.30 | 31.80 | 0.707 |
| inha | 33.17 | 9.78 | 5.82 | 60.87 | 48.83 | 58.14 | 0.769 |
| ital | 20.35 | 4.35 | 2.18 | 32.56 | 21.74 | 21.78 | 0.5 |
| jak2 | 17.76 | 8.04 | 4.95 | 28.78 | 40.18 | 49.50 | 0.775 |
| kif11 | 34.31 | 9.32 | 5.00 | 57.15 | 46.58 | 49.95 | 0.765 |
| kit | 1.81 | 1.20 | 1.75 | 2.83 | 6.00 | 17.50 | 0.663 |
| kith | 51.00 | 15.83 | 8.59 | 100.00 | 78.95 | 85.99 | 0.963 |
| kpcb | 63.90 | 14.23 | 7.70 | 97.72 | 71.08 | 77.00 | 0.888 |
| lck | 11.43 | 4.48 | 3.24 | 17.27 | 22.40 | 32.40 | 0.683 |
| lkha4 | 1.76 | 3.51 | 4.09 | 3.13 | 17.55 | 40.90 | 0.778 |
| mapk2 | 23.95 | 10.51 | 6.14 | 38.72 | 52.47 | 61.40 | 0.854 |
| mcr | 19.30 | 6.81 | 4.68 | 34.63 | 34.05 | 46.80 | 0.779 |
| met | 50.03 | 13.86 | 7.35 | 72.81 | 69.27 | 73.50 | 0.893 |
| mk01 | 31.83 | 13.69 | 7.08 | 54.35 | 68.35 | 70.87 | 0.866 |
| mk10 | 5.77 | 2.12 | 1.35 | 8.95 | 10.59 | 13.50 | 0.537 |
| mk14 | 12.80 | 6.54 | 4.34 | 20.33 | 32.70 | 43.40 | 0.732 |

| Target | $EF^{1\%}$ | $EF^{5\%}$ | $EF^{10\%}$ | $HR^{1\%}$ | $HR^{5\%}$ | $HR^{10\%}$ | AUC |
|---|---|---|---|---|---|---|---|
| mmp13 | 49.30 | 14.96 | 7.76 | 74.80 | 74.80 | 77.60 | 0.912 |
| mp2k1 | 16.47 | 5.29 | 3.88 | 24.10 | 26.42 | 38.80 | 0.730 |
| nos1 | 3.98 | 1.80 | 1.80 | 4.88 | 9.01 | 18.00 | 0.650 |
| nram | 33.66 | 9.38 | 5.71 | 52.38 | 46.92 | 57.10 | 0.862 |
| pa2ga | 7.13 | 5.46 | 4.35 | 13.46 | 27.27 | 43.46 | 0.719 |
| parp1 | 22.87 | 8.54 | 5.61 | 38.04 | 42.70 | 56.10 | 0.845 |
| pde5a | 24.14 | 6.93 | 4.90 | 34.41 | 34.65 | 49.00 | 0.809 |
| pgh1 | 6.15 | 2.56 | 1.95 | 10.91 | 12.81 | 19.50 | 0.624 |
| pgh2 | 29.62 | 10.81 | 6.39 | 54.67 | 54.02 | 63.90 | 0.833 |
| plk1 | 2.81 | 0.94 | 0.94 | 4.36 | 4.70 | 9.39 | 0.5 |
| pnph | 62.69 | 19.79 | 9.91 | 91.54 | 99.05 | 99.10 | 0.996 |
| ppara | 17.72 | 7.30 | 4.93 | 33.50 | 36.48 | 49.30 | 0.793 |
| ppard | 6.23 | 5.17 | 4.04 | 12.00 | 25.85 | 40.40 | 0.77 |
| pparg | 30.42 | 10.29 | 6.12 | 57.20 | 51.45 | 61.20 | 0.826 |
| prgr | 7.18 | 3.14 | 1.88 | 13.20 | 15.71 | 18.80 | 0.633 |
| ptn1 | 21.46 | 5.53 | 3.54 | 37.83 | 27.68 | 35.40 | 0.728 |
| pur2 | 46.75 | 19.23 | 9.81 | 85.19 | 96.01 | 98.00 | 0.992 |
| pygm | 0.00 | 0.26 | 0.78 | 0.00 | 1.30 | 7.81 | 0.5822 |
| pyrd | 46.54 | 10.99 | 6.21 | 78.79 | 54.98 | 62.10 | 0.859 |
| reni | 11.47 | 5.58 | 3.17 | 16.90 | 27.90 | 31.70 | 0.656 |
| rock1 | 9.00 | 5.40 | 4.30 | 14.07 | 27.01 | 43.00 | 0.766 |
| rxra | 5.32 | 2.90 | 1.68 | 9.86 | 14.49 | 16.82 | 0.587 |
| sahh | 55.76 | 19.96 | 10.01 | 100.00 | 100.00 | 100.00 | 1.0 |
| src | 11.44 | 4.05 | 2.73 | 17.14 | 20.25 | 27.30 | 0.665 |
| tgfr1 | 12.07 | 4.81 | 3.31 | 18.60 | 24.07 | 33.10 | 0.781 |
| thb | 26.37 | 9.52 | 5.25 | 36.00 | 47.58 | 52.45 | 0.772 |
| thrb | 2.60 | 2.13 | 1.63 | 4.37 | 10.64 | 16.30 | 0.617 |
| try1 | 18.46 | 6.28 | 3.88 | 31.44 | 31.40 | 38.80 | 0.666 |
| tryb1 | 11.47 | 6.21 | 4.46 | 21.79 | 31.08 | 44.60 | 0.787 |
| tysy | 48.96 | 13.22 | 7.61 | 77.94 | 66.03 | 76.10 | 0.905 |
| urok | 25.93 | 8.52 | 5.12 | 41.99 | 42.58 | 51.20 | 0.783 |
| vgfr2 | 9.06 | 4.69 | 3.47 | 14.62 | 23.45 | 34.70 | 0.675 |
| wee1 | 60.29 | 18.46 | 9.61 | 98.40 | 92.16 | 96.10 | 0.99 |
| xiap | 51.44 | 17.82 | 9.01 | 98.07 | 89.01 | 90.01 | 0.972 |
| Average | 22.28 | 7.63 | 4.77 | 35.95 | 38.37 | 47.90 | 0.77 |

Tab. S2: List of detailed results for InterLig on the standard DUD-E dataset.

| Target | $EF^{1\%}$ | $EF^{5\%}$ | $EF^{10\%}$ | $HR^{1\%}$ | $HR^{5\%}$ | $HR^{10\%}$ | AUC |
|---|---|---|---|---|---|---|---|
| aa2ar | 40.08 | 10.37 | 5.81 | 60.50 | 51.85 | 58.10 | 0.779 |
| abl1 | 17.62 | 6.16 | 4.23 | 29.36 | 30.78 | 42.30 | 0.737 |
| ace | 34.36 | 11.04 | 6.72 | 55.54 | 55.23 | 67.20 | 0.892 |
| aces | 32.64 | 8.70 | 4.94 | 55.43 | 43.50 | 49.40 | 0.715 |
| ada | 36.83 | 10.75 | 6.56 | 61.82 | 53.75 | 65.60 | 0.848 |
| ada17 | 16.43 | 9.50 | 5.53 | 22.59 | 47.50 | 55.30 | 0.85 |
| adrb1 | 18.60 | 10.12 | 5.87 | 28.56 | 50.60 | 58.70 | 0.842 |
| adrb2 | 3.47 | 1.82 | 1.86 | 5.27 | 9.10 | 18.60 | 0.668 |
| akt1 | 10.93 | 6.28 | 4.51 | 19.17 | 31.42 | 45.10 | 0.792 |

| Target | $EF^{1\%}$ | $EF^{5\%}$ | $EF^{10\%}$ | $HR^{1\%}$ | $HR^{5\%}$ | $HR^{10\%}$ | AUC |
|---|---|---|---|---|---|---|---|
| akt2 | 16.26 | 5.48 | 3.50 | 27.15 | 27.37 | 35.00 | 0.794 |
| aldr | 32.24 | 10.45 | 6.10 | 56.04 | 52.20 | 61.00 | 0.853 |
| ampc | 26.90 | 6.67 | 3.54 | 44.83 | 33.35 | 35.40 | 0.724 |
| andr | 26.39 | 6.69 | 4.39 | 48.63 | 33.45 | 43.90 | 0.768 |
| aofb | 2.47 | 1.31 | 1.23 | 4.29 | 6.55 | 12.30 | 0.626 |
| bace1 | 7.10 | 2.84 | 2.13 | 9.84 | 14.19 | 21.30 | 0.685 |
| braf | 21.03 | 8.43 | 5.07 | 31.68 | 42.11 | 50.70 | 0.78 |
| cah2 | 0.41 | 0.73 | 1.16 | 0.64 | 3.65 | 11.60 | 0.811 |
| casp3 | 5.52 | 3.82 | 2.91 | 10.09 | 19.09 | 29.10 | 0.611 |
| cdk2 | 6.76 | 2.87 | 2.11 | 11.30 | 14.35 | 21.10 | 0.612 |
| comt | 38.91 | 16.13 | 9.02 | 41.02 | 80.45 | 90.20 | 0.974 |
| cp2c9 | 0.83 | 0.67 | 1.25 | 1.32 | 3.35 | 12.50 | 0.543 |
| cp3a4 | 8.79 | 3.29 | 2.41 | 12.50 | 16.45 | 24.10 | 0.648 |
| csf1r | 28.91 | 10.00 | 6.14 | 39.02 | 50.00 | 61.40 | 0.861 |
| cxcr4 | 42.11 | 9.43 | 6.10 | 44.12 | 47.20 | 61.06 | 0.858 |
| def | 46.24 | 13.24 | 7.37 | 75.87 | 66.30 | 73.70 | 0.898 |
| dhi1 | 14.21 | 5.03 | 3.33 | 23.86 | 25.15 | 33.30 | 0.667 |
| dpp4 | 20.31 | 6.32 | 3.99 | 26.09 | 31.60 | 39.90 | 0.754 |
| drd3 | 6.04 | 1.63 | 1.19 | 8.41 | 8.15 | 11.90 | 0.578 |
| dyr | 26.93 | 12.78 | 7.52 | 35.64 | 63.90 | 75.20 | 0.916 |
| egfr | 34.36 | 12.40 | 7.21 | 52.39 | 62.00 | 72.10 | 0.893 |
| esr1 | 46.64 | 13.14 | 7.62 | 84.75 | 65.70 | 76.20 | 0.896 |
| esr2 | 27.38 | 10.61 | 6.46 | 48.30 | 53.05 | 64.60 | 0.863 |
| fa10 | 9.70 | 4.21 | 2.81 | 25.35 | 21.05 | 28.10 | 0.709 |
| fa7 | 0.00 | 10.72 | 7.19 | 0.00 | 53.52 | 71.90 | 0.924 |
| fabp4 | 33.95 | 7.64 | 4.26 | 57.15 | 38.31 | 42.56 | 0.789 |
| fak1 | 43.40 | 17.63 | 9.20 | 79.63 | 87.97 | 92.00 | 0.969 |
| fgfr1 | 0.68 | 0.71 | 1.08 | 20.00 | 20.88 | 31.76 | 0.452 |
| fkb1a | 14.43 | 6.13 | 4.32 | 27.11 | 30.60 | 43.20 | 0.811 |
| fnta | 21.71 | 8.07 | 5.59 | 24.22 | 40.35 | 55.90 | 0.821 |
| fpps | 82.42 | 19.54 | 10.00 | 82.35 | 97.60 | 100.00 | 0.997 |
| gcr | 29.52 | 8.52 | 4.89 | 49.99 | 42.62 | 48.90 | 0.774 |
| glcm | 0.00 | 1.85 | 1.67 | 0.00 | 9.27 | 16.68 | 0.665 |
| gria2 | 39.83 | 10.76 | 6.20 | 52.50 | 53.77 | 62.00 | 0.816 |
| grik1 | 12.96 | 4.76 | 3.86 | 19.69 | 23.78 | 38.60 | 0.696 |
| hdac2 | 15.84 | 4.47 | 2.82 | 25.72 | 22.33 | 28.20 | 0.672 |
| hdac8 | 5.85 | 3.51 | 3.83 | 8.50 | 17.55 | 38.30 | 0.772 |
| hivint | 1.01 | 3.60 | 2.90 | 1.50 | 18.00 | 29.00 | 0.76 |
| hivpr | 31.53 | 12.23 | 6.92 | 44.19 | 61.15 | 69.20 | 0.878 |
| hivrt | 10.06 | 4.91 | 3.46 | 17.71 | 24.55 | 34.60 | 0.703 |
| hmdh | 47.12 | 13.99 | 7.47 | 89.89 | 70.02 | 74.70 | 0.889 |
| hs90a | 16.48 | 7.93 | 5.76 | 24.49 | 39.71 | 57.60 | 0.869 |
| hxk4 | 33.61 | 11.76 | 6.19 | 64.58 | 58.71 | 61.90 | 0.894 |
| igf1r | 21.02 | 7.83 | 5.75 | 32.98 | 39.17 | 57.50 | 0.802 |
| inha | 28.40 | 6.51 | 3.72 | 52.17 | 32.53 | 37.20 | 0.784 |
| ital | 18.88 | 4.49 | 2.39 | 30.23 | 22.45 | 23.90 | 0.501 |
| jak2 | 17.74 | 7.84 | 4.77 | 28.78 | 39.24 | 47.65 | 0.801 |
| kif11 | 40.26 | 9.30 | 5.43 | 67.14 | 46.52 | 54.35 | 0.732 |
| kit | 2.41 | 0.96 | 0.90 | 3.77 | 4.80 | 9.00 | 0.567 |

| Target | $EF^{1\%}$ | $EF^{5\%}$ | $EF^{10\%}$ | $HR^{1\%}$ | $HR^{5\%}$ | $HR^{10\%}$ | AUC |
|---|---|---|---|---|---|---|---|
| kith | 50.98 | 14.77 | 7.88 | 100.00 | 73.70 | 78.88 | 0.952 |
| kpcb | 63.46 | 13.67 | 7.48 | 88.64 | 68.32 | 74.80 | 0.89 |
| lck | 8.37 | 4.11 | 3.04 | 12.60 | 20.55 | 30.40 | 0.639 |
| lkha4 | 11.13 | 6.20 | 4.44 | 19.79 | 31.00 | 44.40 | 0.808 |
| mapk2 | 25.19 | 11.01 | 6.50 | 40.33 | 55.00 | 65.07 | 0.875 |
| mcr | 16.06 | 8.50 | 5.00 | 28.85 | 42.56 | 49.95 | 0.803 |
| met | 32.52 | 14.70 | 8.01 | 47.36 | 73.50 | 80.10 | 0.905 |
| mk01 | 41.95 | 12.66 | 6.84 | 71.73 | 63.30 | 68.40 | 0.887 |
| mk10 | 4.81 | 1.35 | 1.06 | 7.46 | 6.75 | 10.60 | 0.477 |
| mk14 | 15.21 | 6.89 | 4.45 | 24.17 | 34.45 | 44.50 | 0.74 |
| mmp13 | 44.70 | 15.69 | 8.39 | 67.65 | 78.45 | 83.90 | 0.931 |
| mp2k1 | 13.16 | 3.64 | 3.06 | 19.27 | 18.19 | 30.60 | 0.688 |
| nos1 | 2.01 | 2.20 | 1.80 | 2.47 | 11.00 | 18.00 | 0.653 |
| nram | 34.68 | 8.36 | 4.90 | 53.97 | 41.82 | 49.00 | 0.832 |
| pa2ga | 11.21 | 5.06 | 3.44 | 21.16 | 25.27 | 34.37 | 0.688 |
| parp1 | 25.81 | 8.58 | 6.02 | 42.95 | 42.92 | 60.20 | 0.876 |
| pde5a | 23.12 | 5.68 | 3.97 | 32.97 | 28.39 | 39.70 | 0.784 |
| pgh1 | 4.60 | 2.57 | 1.90 | 8.18 | 12.84 | 19.00 | 0.612 |
| pgh2 | 29.78 | 11.11 | 6.91 | 54.89 | 55.52 | 69.10 | 0.829 |
| plk1 | 3.77 | 1.51 | 1.51 | 5.79 | 7.55 | 15.10 | 0.493 |
| pnph | 61.72 | 19.78 | 9.91 | 90.14 | 99.00 | 99.10 | 0.996 |
| ppara | 17.69 | 7.67 | 5.09 | 33.49 | 38.35 | 50.90 | 0.81 |
| ppard | 14.22 | 8.26 | 5.83 | 27.41 | 41.28 | 58.30 | 0.863 |
| pparg | 35.13 | 10.75 | 6.53 | 66.15 | 53.75 | 65.30 | 0.865 |
| prgr | 6.15 | 3.07 | 2.29 | 11.33 | 15.36 | 22.90 | 0.599 |
| ptn1 | 22.19 | 5.85 | 3.62 | 39.18 | 29.24 | 36.20 | 0.677 |
| pur2 | 54.88 | 20.03 | 10.01 | 100.00 | 100.00 | 100.00 | 1.0 |
| pygm | 2.61 | 3.37 | 2.21 | 5.01 | 16.87 | 22.12 | 0.621 |
| pyrd | 47.41 | 10.80 | 6.13 | 80.29 | 54.05 | 61.24 | 0.854 |
| reni | 0.00 | 9.51 | 5.24 | 0.00 | 47.57 | 52.40 | 0.738 |
| rock1 | 11.98 | 5.19 | 4.20 | 18.75 | 25.99 | 42.00 | 0.74 |
| rxra | 0.00 | 1.07 | 0.92 | 0.00 | 5.35 | 9.20 | 0.520 |
| sahh | 55.71 | 19.94 | 10.00 | 100.00 | 100.00 | 100.00 | 1.0 |
| src | 8.95 | 3.78 | 3.04 | 13.14 | 18.90 | 30.40 | 0.694 |
| tgfr1 | 12.07 | 4.21 | 4.28 | 18.61 | 21.04 | 42.80 | 0.807 |
| thb | 33.10 | 9.90 | 5.63 | 45.33 | 49.50 | 56.30 | 0.758 |
| thrb | 2.02 | 2.93 | 2.39 | 3.28 | 14.66 | 23.90 | 0.678 |
| try1 | 18.31 | 6.77 | 4.28 | 30.43 | 33.85 | 42.80 | 0.718 |
| tryb1 | 12.13 | 8.51 | 5.94 | 23.08 | 42.57 | 59.40 | 0.85 |
| tysy | 55.40 | 14.69 | 8.08 | 88.23 | 73.41 | 80.72 | 0.925 |
| urok | 25.90 | 7.65 | 4.82 | 42.00 | 38.29 | 48.20 | 0.802 |
| vgfr2 | 11.74 | 5.23 | 4.33 | 18.97 | 26.16 | 43.30 | 0.755 |
| wee1 | 60.14 | 19.00 | 9.70 | 98.38 | 95.10 | 97.10 | 0.99 |
| xiap | 51.43 | 18.81 | 9.41 | 98.07 | 93.96 | 94.01 | 0.97 |
| Average | 23.13 | 8.07 | 4.91 | 37.32 | 40.54 | 49.38 | 0.78 |

Tab. S3: List of detailed results for InterLig on the multiple conformers DUD-E dataset.

| | $EF^{1\%}$ | $EF^{5\%}$ | $EF^{10\%}$ | AUC |
|---|---|---|---|---|
| LS-align rigid + OMEGA | 22.77 | 7.5 | 4.63 | 0.75 |
| LS-align flexi | 21.57 | 7.34 | 4.54 | 0.75 |
| InterLig | **23.13** | **8.07** | **4.91** | **0.78** |

Tab. S4: Comparison of performance between LS-align rigid on the set of conformers generated with OMEGA and LS-align run on the regular set in "flexible" mode, where LS-align generates its own conformers. It is actually slightly better to run LS-align in the default "rigid" mode on the set of conformers generated by OMEGA. This differs from the experiments in the original LS-align paper, where LS-align performed more or less equally in "flexible" mode and in "rigid" mode on a set of conformers generated with OMEGA. This is most likely because of a difference of parameters used to run OMEGA (a fixed number of conformers was generated for each molecule in the LS-align paper, while we generated a variable number of conformers with the constraint that no two conformers can be closer than 2ÅRMSD). Nevertheless, InterLig performs better than both versions of LS-align.

| Target | N. decoys | Memory | Time | Seed size | Avg. decoy size |
|---|---|---|---|---|---|
| dhi1 | 90,259 | 0.9GB | 15min | 29 | 27.6 |
| adrb2 | 143,442 | 1.6GB | 17min | 20 | 31.5 |
| fnta | 375,425 | 4.3GB | 105min | 43 | 31.9 |

Tab. S5: Execution time and memory requirements to run InterLig on three sets of decoys generated with OMEGA. The memory scales linearly with the size of the set, while the execution time is also dependent on the size of the seed ligand and decoys. The tests were run on a single core of an Ubuntu 14.04 with a 3.4GHz CPU and 32GB RAM.

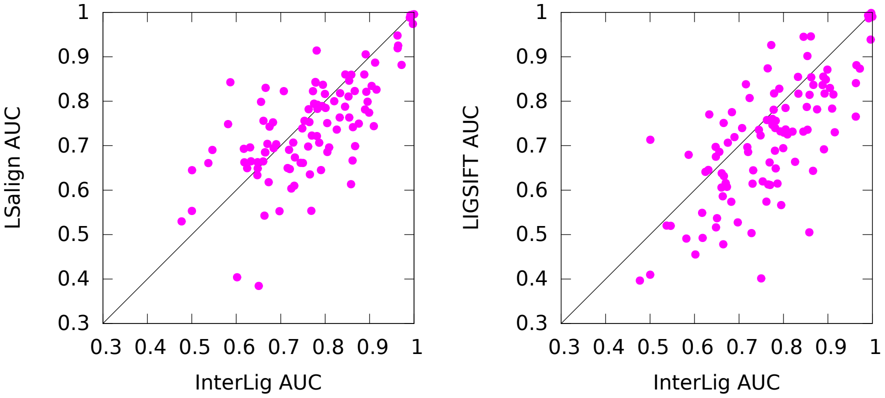


(a)

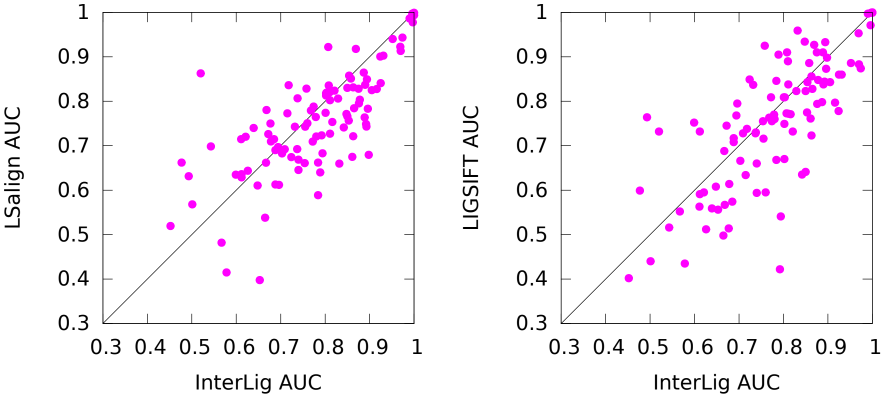


(b)

Fig. S3: Performance comparison of InterLig against LS-align (left side) and LIGSIFT (right side). Plot (a) compares the AUC for the 102 DUD-E targets on the regular set of ligands and decoys, plot (b) compares the AUC for the 102 DUD-E targets on the multiple conformer set of DUD-E ligands and decoys.

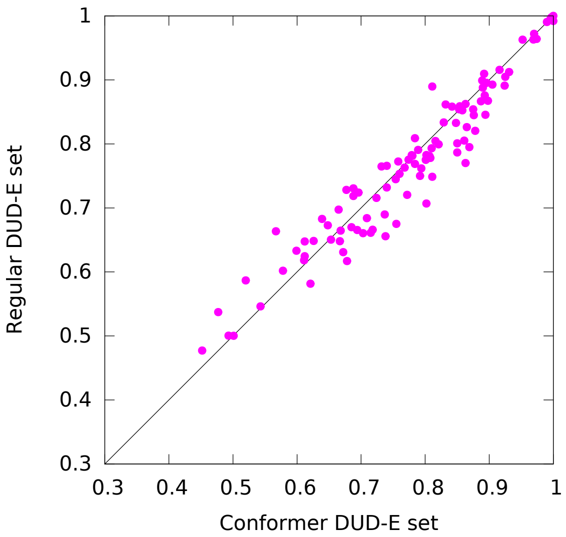


Fig. S4: Comparison of InterLig AUC for 102 DUD-E targets on the regular set of ligands and decoys versus the multiple conformers set. From this experiment, it appears that it is more helpful to have conformers in cases where InterLig already performs well (higher AUC) on the regular set. Results do not improve with conformers when the AUC on the regular set is marginal (closer to random, 0.5).

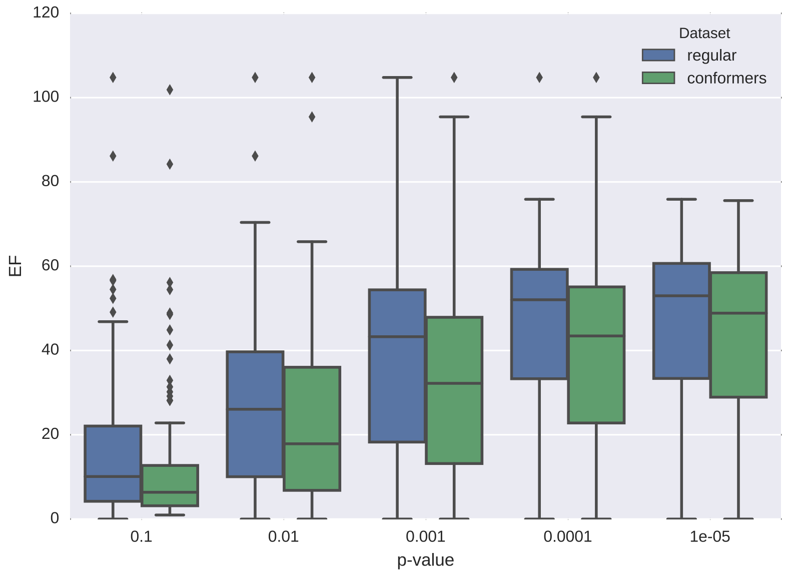


Fig. S5: Boxplot of the enrichment values when selecting the top alignments with a p-value threshold rather than a fixed top percentage of the ranked list. It is no surprise that the median enrichment is lower in the conformers case, since using conformers will yield on average lower p-values (more experiments run on the same set). So stricter p-value thresholds are needed when working with conformer sets.
